# Diving beetle offspring oviposited in amphibian spawn prey on the tadpoles upon hatching

**DOI:** 10.1101/666008

**Authors:** John Gould, Jose W. Valdez, Simon Clulow, John Clulow

## Abstract

In highly ephemeral freshwater habitats, predatory vertebrates are typically unable to become established, leaving an open niche often filled by macroinvertebrate predators. However, these predators are faced with the challenge of finding sufficient food sources as the rapid rate of desiccation prevents the establishment of extended food chains and limits the number of prey species present. It could therefore be advantageous for adults to oviposit their offspring in the presence of future prey within sites of extreme ephemerality. We report the first case of adult diving beetles ovipositing their eggs within spawn of the sandpaper frog, *Lechriodus fletcheri*. This behaviour was found among several pools used by *L. fletcheri* for reproduction. Beetle eggs oviposited in frog spawn were found to hatch within 24 hours of the surrounding *L. fletcheri* eggs, with the larvae becoming voracious consumers of the hatched tadpoles. Although it has yet to be established experimentally whether this is an adaptive behaviour, the laying of eggs among potential future tadpole prey in this instance should confer significant fitness benefits for the offspring upon hatching, ensuring that they are provided an immediate source of food at the start of their development and potentially throughout. This oviposition behaviour may be common among diving beetles and could form a significant predatory threat for amphibians with a free-swimming larval stage in ephemeral freshwater habitats.

## Introduction

Diving beetles (Coleoptera: Dytiscidae) are a large invertebrate group comprising more than 4,000 species which exploit a variety of freshwater aquatic habitats for reproduction, development, and feeding (Larson, Alarie et al. 2000, Yee 2014). Despite their ubiquity, they remain understudied as an aquatic insect group, with many aspects of their ecology still relatively unknown (Yee 2014). While adult forms are considered scavengers, their larvae are exclusively carnivorous and are often the top predator in freshwater habitats when vertebrate predators, such as fish, are absent (Wellborn, Skelly et al. 1996, Culler, Ohba et al. 2014). Upon hatching, Dytiscidae larvae are equipped with a variety of adaptations that make them effective hunters, including sensory organs for perceiving tactile and chemical cues (Culler, Ohba et al. 2014), as well as visual scanning behaviours (Formanowicz 1987, Buschbeck, Sbita et al. 2007) that allow prey to be quickly detected. Once a potential prey item has been located, larvae will capture them using their large piercing mandibles (Kruse 1983, Formanowicz 1987, Morgan 1992), which are then used to inject digestive enzymes into the prey so that the liquidised remains can be sucked up and ingested (Kruse 1983, Kehl 2014). A variety of tactics are used by larvae to catch prey, such as sit-and-wait ambush or search-and-follow hunting, with some species having the capacity to shift tactics based on surrounding conditions (e.g. prey density, habitat complexity) (Formanowicz 1987, Michel and Adams 2009, Yee 2010, Yee 2014).

Diving beetles are central to freshwater food webs, with the ability to cause trophic cascades due to their high rate of predation on a diverse number of aquatic invertebrates and smaller vertebrates (Cobbaert, Bayley et al. 2010, Culler, Ohba et al. 2014). Their voracious appetite also includes the tadpoles of several amphibian species (Brodie, Formanowicz et al. 1978, Formanowicz and Bobka 1989, Tejedo 1993, Valdez 2019), with experiments showing larvae capable of consuming dozens of tadpoles in a single day (Kruse 1983). They are such effective predators that tadpoles may undergo dytiscid-induced behavioural changes such as altering levels of activity (Tejedo 1993), avoiding areas where larvae are present (Rubbo, Mirza et al. 2006, Smith and Awan 2009), and even shortening hatching time or reducing the size of tadpoles when compared to sites where they are not present (Pearman 1995).

Since diving beetles are common and abundant within many aquatic habitats (Gioria 2014), it is likely that the presence of larvae will coincide with the developmental period of many amphibians whose life history includes a free-swimming tadpole stage (Duellman 1992, Heard, Scroggie et al. 2012). As adult beetles are one of the first macroinvertebrates to colonize newly formed aquatic habitats (Lundkvist, Landin et al. 2003, Bilton 2014), their subsequent offspring may be a significant source of predation of tadpoles of amphibians that also exploit ephemeral habitats for reproduction (Gill 1978, Marsh and Trenham 2001). However, the predation threat diving beetles pose to amphibian larvae is rarely observed in nature, with many aspects of their interactions, egg deposition behaviour, and larval development still relatively unknown (Kehl 2014, Yee 2014). Herein, we report the first documented case of diving beetle ovipositing their eggs within amphibian spawn (sandpaper frog, *Lechriodus fletcheri*), with both larvae and amphibian eggs hatching at similar times. We also describe the predatory behaviour of one species of diving beetle larvae on the recently hatched tadpoles.

## Observations

A recently oviposited *L. fletcheri* spawn was found on December 2, 2013 in a small pool on the forest floor within the Watagan Mountain Range in New South Wales, Australia (33°0’18.396” S, 151°26’ 23.4312” E) during an ecological study on the species (data not presented here). Similar pools are found throughout the Watagan Mountains and are often highly ephemeral, drying out within days or weeks after rainfall has ceased. They are also almost exclusively colonised by the sandpaper frog *L. fletcheri* and/or mosquito larvae (*pers. obs*.). However, the spawn in this particular pool was found to contain several unidentified eggs interspersed through a small section of the frothed oviduct fluid that surrounds the residing embryos throughout embryogenesis. The unidentified eggs were between 3-4 mm and approximately half the length of the neighbouring amphibian embryos. The spawn with unidentified eggs was taken back to the laboratory at the University of Newcastle and placed in a small container filled with aged tap water for the eggs to continue developing. The eggs were maintained at room temperature, approximately 23°C. Upon hatching, the smaller eggs were later identified as a Dytiscidae species by the unique morphology of the offspring, including an elongated body and large curved mandibles. As it was not known at the time that the beetle larvae were a predation threat to the hatched tadpoles, they were left to develop in the containers until predation was observed, after which the beetle larvae were removed.

An additional 65 recently laid spawn collected from 16 pools between December 2013 and February 2014 were also inspected for the presence of diving beetle eggs, thereby allowing us to determine whether they are regularly laid in *L. fletcheri* spawn. The number of beetle eggs laid per spawn were recorded when present. Each spawn was kept in a separate container filled with aged tap water and allowed to develop normally, being checked daily to determine relative hatching times of the beetle and amphibian eggs. Over this same period, pools within a localised area of the Watagan Mountains were surveyed for the presence of adult diving beetles. A breeding pair still in amplexus was found and brought back to the laboratory, where they were placed in a small container filled with aged tap water and exposed to a recently laid *L. fletcheri* spawn devoid of any diving beetle eggs, to determine whether they would lay in the spawn. The spawn was checked daily to determine the timing of hatching of the amphibian and diving beetle eggs.

All diving beetle eggs, including the adult pair that was collected, were found to belong to the same species of diving beetle (*Hydaticus parallelus*; Figs 1 & 2). This was the only species of diving beetle present across the pools surveyed. Diving beetle eggs were found in 7 of the 65 spawns (11%) from 3 different pools, while adults and larvae of the beetle were observed across several pools. The number of unhatched diving beetle eggs present within the spawns ranged between 1 and 175 (median=7) and nearly all hatched within 24 hours of the neighbouring amphibian eggs hatching, which typically occurred between 48 and 72 hours after the spawn was collected. Upon hatching, the beetle larvae were similar in size to the hatched tadpoles. Larvae were frequently observed to adopt an immobile stance near the surface of the water, inverted with their heads facing downwards and mandibles open. After adopting this immobile stance, larvae were often observed attacking tadpoles that approached. On some occasions, beetle larvae would swim towards the bottom of the container where the tadpoles were residing and pursue nearby tadpoles in an attempt to catch them between opened mandibles. The beetle larvae were observed to catch multiple tadpoles within a matter of hours. Often, beetle larvae would partially consume the tadpole they had caught before mounting an attack on another tadpole that swam close by.

**Figure 1.**
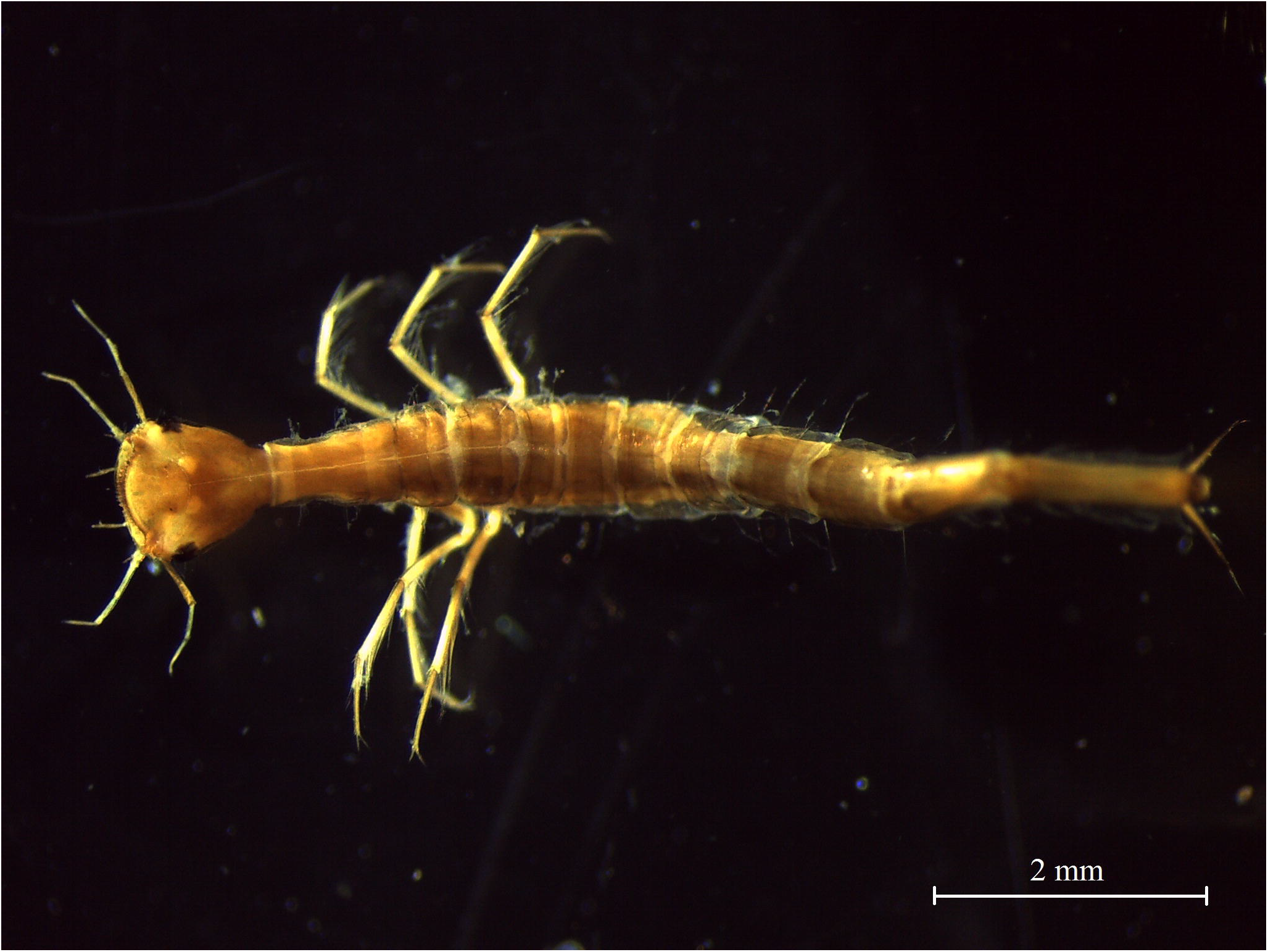
*Hydaticus parallelus* larvae a few days post-hatching, revealing the elongated, segmented body plan shared by all diving beetle larvae.

**Figure 2.**
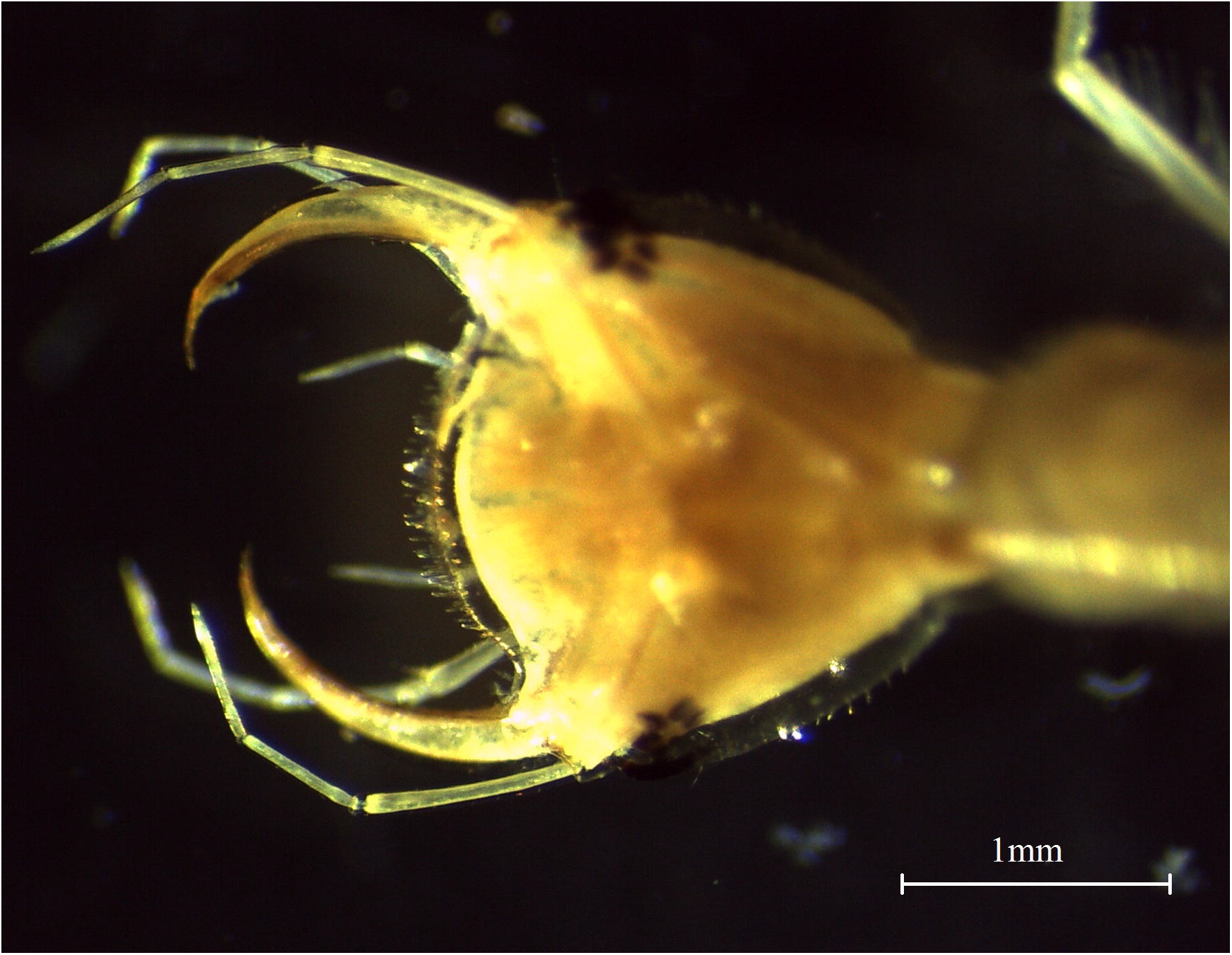
Close-up of the head of *Hydaticus parallelus* larvae, showing the opened mandibles.

The adult beetle pair that was collected remained in amplexus for several days, laying between 3-5 eggs within the frog spawn that was present within their container (Fig. 3). Approximately 3 days later, tadpoles hatched from the spawn body, with the diving beetle eggs all hatching less than 24 hours afterwards.

**Figure 3.**
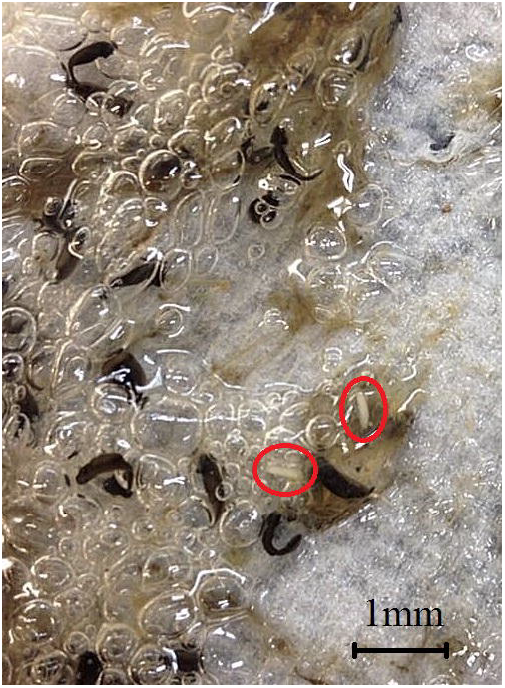
*Lechriodus fletcheri* spawn with several *Hydaticus parallelus* eggs (red circles) interspersed through the frothed oviduct fluid of the spawn body.

## Discussion

We determined that *H. parallelus* adults oviposit their eggs within *L. fletcheri* spawn and suggest they time this so that both eggs types hatch together in order to allow the beetle larvae to prey upon the newly hatched tadpoles. As far as we are aware, this is the first documented case of this ovipositioning behaviour among diving beetles. Adult diving beetles are often the first macroinvertebrate predator to colonise newly formed ephemeral habitats (Lundkvist, Landin et al. 2003) which occurs at a similar time to that of *L. fletcheri*, which also oviposit their eggs in those habitats (Clulow and Swan 2018). Therefore, we suggest the observed behaviour is likely adaptive and that the tadpoles are a primary food source for diving beetle larvae in these habitats. *Hydaticus parallelus* larvae appear to be the apex predator within aquatic sites used by *L. fletcheri* for reproduction, given that the extreme ephemerality of these sites prevent other common aquatic predators from colonising (Skelly 1996, Wellborn, Skelly et al. 1996), and no other predator was found during these surveys. As such, they may constitute a significant source of pre-metamorphic mortality for *L. fletcheri* alongside pool desiccation and cannibalism (Clulow & Swan, 2018), although these latter two sources likely still constitute the majority of *L. fletcheri* mortalities in these habitats.

It remains unclear whether adult beetles are actively seeking out frog spawn to oviposit within or whether it occurs opportunistically through chance encounters as a convenient substrate to deposit eggs. However, considering that oviposition in the spawn allows for the beetle offspring to obtain a readily accessible, high-protein feed, it is likely there is selection towards this behaviour. This may be highly beneficial, considering that the only alternative food source that regularly occurs at these sites that we are aware of are mosquito larvae, which are a comparatively smaller prey item that will be less nutritious per catch. It also remains unproven whether the beetle eggs are deliberately laid at a time to enable synchronisation of hatching with their tadpole counterparts, or possibly even if the process of tadpole hatching may be an environmentally-cued hatching trigger that causes the beetles to hatch (Warkentin 2011, Doody and Paull 2013). Previous evidence suggests diving beetle eggs hatch 5-22 days after being oviposited depending on water temperature (Aiken 1986, Inoda 2003). Here, we found hatching times between 4-5 days across all infected spawn examined. We therefore raise the interesting possibility that the laying of the beetle eggs are timed to hatch at similar times to the tadpole eggs, either through timing of oviposition or through environmentally-cued hatching.

It appears that diving beetle larvae prey on *L. fletcheri* tadpoles using both sit-and-wait and swim-and-hunt predation techniques that have previously been reported for species in this group (Michel and Adams 2009, Yee 2010, Yee 2014). As the aquatic sites used by *L fletcheri* are small pools that often lack submerged vegetation, the tactile and chemical cues used by beetle larvae to sense the presence of nearby prey are likely to be highly effective (Michel and Adams 2009, Yee 2010, Yee 2014) given that there is effectively no structural complexity to obstruct the transport of these cues (Formanowicz 1987). The lack of submerged vegetation also reduces available refuges for tadpoles to evade this predation threat, besides dead *Eucalyptus* leaves that periodically fall to the bottom of these pools (pers. obs.). Although it remains undetermined whether *L. fletcheri* tadpoles have developed any specific defences to minimise predation by diving beetle larvae, other tadpole species are known to reduce their activity levels in their presence (Tejedo 1993) to avoid detection or reduce encounter rates, while other species hasten embryogenesis to avoid predation (Johnson, Saenz et al. 2003).

The synchronisation of behaviour between con- and hetero-specifics represents an important interaction between individuals that may confer significant benefits to some or all involved (Evans and Patterson 1971, Soong and Cho 1998, Dostálková and Špinka 2007, Silva, Prieto et al. 2013). Our observations allude to the synchronisation of *H. parallelus* reproductive activity with that of *L. fletcheri*. By ovipositing within frog spawn, adult diving beetles may be improving the fitness of their offspring by placing them within the vicinity of a nutritious food source that can be exploited throughout development. As such, while it is likely to be highly beneficial for diving beetle offspring, it will be highly detrimental for *L. fletcheri* offspring as they will be exposed to a voracious predator type the moment they hatch. Further research will be required to determine whether adults are choosing *L. fletcheri* spawn as an ovipositioning site for an adaptive purpose, or whether spawn merely represents an alternative pool structure that can be used for egg attachment. Given the common occurrence of diving beetle larvae across various freshwater habitats, these results suggest they may be an important controlling mechanism regulating population size for other amphibian species with a free-swimming larval stage.

## Acknowledgements

The authors thank Chris Watt for assisting in species identification of diving beetle specimens. All work was conducted under approval from the University of Newcastle’s Animal Care and Ethics Committee (project no. A-2011-138).

## References

Aiken, R. B. (1986). “Effects of Temperature on Incubation Times and Mortality Rates of Eggs of *Dytiscus alaskanus* (Coleoptera: Dytiscidae).” Holarctic Ecology 9(2): 133–136.

Bilton, D. T. (2014). Dispersal in Dytiscidae. Ecology, Systematics, and the Natural History of Predaceous Diving Beetles (Coleoptera: Dytiscidae). D. A. Yee. Dordrecht, Springer Netherlands: 387–407.

Brodie, E. D., D. R. Formanowicz and E. Brodie III (1978). “The development of noxiousness of *Bufo americanus* tadpoles to aquatic insect predators.” Herpetologica 34(3): 302–306.

Buschbeck, E. K., S. J. Sbita and R. C. Morgan (2007). “Scanning behavior by larvae of the predacious diving beetle, Thermonectus marmoratus (Coleoptera: Dytiscidae) enlarges visual field prior to prey capture.” Journal of Comparative Physiology A 193(9): 973–982.

Clulow, S. and M. Swan (2018). “A complete guide to frogs of Australia.”

Cobbaert, D., S. E. Bayley and J.-L. Greter (2010). "Effects of a top invertebrate predator (*Dytiscus alaskanus*; Coleoptera: Dytiscidae) on fishless pond ecosystems.” Hydrobiologia 644(1): 103–114.

Culler, L. E., S. Ohba and P. Crumrine (2014). Predator-prey interactions of dytiscids. Ecology, Systematics, and the Natural History of Predaceous Diving Beetles (Coleoptera: Dytiscidae). D. A. Yee. Dordrecht, Springer Netherlands: 363–386.

Doody, J. S. and P. Paull (2013). "Hitting the ground running: environmentally cued hatching in a lizard.” Copeia 2013(1): 160–165.

Dostálková, I. and M. Špinka (2007). "Synchronization of behaviour in pairs: the role of communication and consequences in timing.” Animal Behaviour 74(6): 1735–1742.

Duellman, W. E. (1992). "Reproductive strategies of frogs.”. Scientific American 267(1): 80–87.

Evans, S. and G. Patterson (1971). "The synchronization of behaviour in flocks of estrildine finches.” Animal Behaviour 19(3): 429–438.

Formanowicz, D. R. (1987). “Foraging tactics of *Dytiscus verticalis* larvae (Coleoptera: Dytiscidae): prey detection, reactive distance and predator size.” Journal of the Kansas Entomological Society: 92–99.

Formanowicz, D. R. and M. S. Bobka (1989). "Predation risk and microhabitat preference: an experimental study of the behavioral responses of prey and predator.”. American Midland Naturalist: 379–386.

Gill, D. E. (1978). “The metapopulation ecology of the red-spotted newt, Notophthalmus viridescens (*Rafinesque*).” Ecological Monographs 48(2): 145–166.

Gioria, M. (2014). Habitats. Ecology, Systematics, and the Natural History of Predaceous Diving Beetles (Coleoptera: Dytiscidae). D. A. Yee. Dordrecht, Springer Netherlands: 307–362.

Heard, G. W., M. P. Scroggie and B. S. Malone (2012). “Classical metapopulation theory as a useful paradigm for the conservation of an endangered amphibian.”. Biological Conservation 148(1): 156–166.

Inoda, T. (2003). “Mating and reproduction of predaceous diving beetles, *Dytiscus sharpi*, observed under artificial breeding conditions.” Zoological Science 20(3): 377–382.

Johnson, J. B., D. Saenz, C. K. Adams and R. N. Conner (2003). “The influence of predator threat on the timing of a life-history switch point: predator-induced hatching in the southern leopard frog (*Rana sphenocephala*).” Canadian Journal of Zoology 81(9): 1608–1613.

Kehl, S. (2014). Morphology, anatomy, and physiological aspects of dytiscids. Ecology, Systematics, and the Natural History of Predaceous Diving Beetles (Coleoptera: Dytiscidae). D. A. Yee. Dordrecht, Springer Netherlands: 173–198.

Kruse, K. C. (1983). “Optimal foraging by predaceous diving beetle larvae on toad tadpoles.” Oecologia 58(3): 383–388.

Larson, D., Y. Alarie and R. E. Roughley (2000). Predaceous diving beetles (Coleoptera: Dytiscidae) of the Nearctic Region, with emphasis on the fauna of Canada and Alaska, NRC Research Press.

Lundkvist, E., J. Landin, M. Jackson and C. Svensson (2003). “Diving beetles (Dytiscidae) as predators of mosquito larvae (Culicidae) in field experiments and in laboratory tests of prey preference.”. Bulletin of entomological research 93(3): 219–226.

Marsh, D. M. and P. C. Trenham (2001). "Metapopulation dynamics and amphibian conservation.”. Conservation Biology 15(1): 40–49.

Michel, M. J. and M. M. Adams (2009). "Differential effects of structural complexity on predator foraging behavior.” Behavioral Ecology 20(2): 313–317.

Morgan, R. (1992). Natural history, captive management and the display of the sunburst diving beetle Thermonectus marmoratus. Proceedings of the AAZPA Annual Conference.

Pearman, P. B. (1995). "Effects of pond size and consequent predator density on two species of tadpoles.” Oecologia 102(1): 1–8.

Rubbo, M., R. Mirza, L. Belden, J. Falkenbach, S. Storrs and J. Kiesecker (2006). "Evaluating a predator– prey interaction in the field: the interaction between beetle larvae (predator) and tadpoles (prey).” Journal of Zoology 269(1): 1–5.

Silva, M. A., R. Prieto, I. Jonsen, M. F. Baumgartner and R. S. Santos (2013). "North Atlantic blue and fin whales suspend their spring migration to forage in middle latitudes: building up energy reserves for the journey?” PLoS One 8(10): e76507.

Skelly, D. K. (1996). “Pond drying, predators, and the distribution of Pseudacris tadpoles.” Copeia: 599–605.

Smith, G. R. and A. R. Awan (2009). “The roles of predator identity and group size in the antipredator responses of american toad (*Bufo americanus*) and bullfrog (*Rana catesbeiana*) tadpoles.” Behaviour 146(2): 225–243.

Soong, K. and L. Cho (1998). “Synchronized release of medusae from three species of hydrozoan fire corals.” Coral Reefs 17(2): 145–154.

Tejedo, M. (1993). “Size-dependent vulnerability and behavioral responses of tadpoles of two anuran species to beetle larvae predators.” Herpetologica 49(3): 287–294.

Valdez, J. (2019). “Predaceous Diving Beetles (*Coleoptera: Dytiscidae*) May Affect the Success of Amphibian Conservation Efforts.” Preprints 2019030213.

Warkentin, K. M. (2011). “Environmentally cued hatching across taxa: embryos respond to risk and opportunity.” Integrative and Comparative Biology 51(1): 14–25.

Wellborn, G. A., D. K. Skelly and E. E. Werner (1996). “Mechanisms creating community structure across a freshwater habitat gradient.” Annual review of ecology and systematics 27(1): 337–363.

Yee, D. A. (2010). “Behavior and aquatic plants as factors affecting predation by three species of larval predaceous diving beetles (Coleoptera: Dytiscidae).” Hydrobiologia 637(1): 33–43.

Yee, D. A. (2014). Ecology, Systematics, and the Natural History of Predaceous Diving Beetles (Coleoptera: Dytiscidae). Dordrecht, Springer Netherlands.

